# *linus*: Conveniently explore, share, and present large-scale biological trajectory data from a web browser

**DOI:** 10.1101/2020.04.17.043323

**Authors:** Johannes Waschke, Mario Hlawitschka, Kerim Anlas, Vikas Trivedi, Ingo Roeder, Jan Huisken, Nico Scherf

## Abstract

In biology, we are often confronted with information-rich, large-scale trajectory data, but exploring and communicating patterns in such data is often a cumbersome task. Ideally, the data should be wrapped with an interactive visualisation in one concise package that makes it straightforward to create and test hypotheses collaboratively. To address these challenges, we have developed a tool, *linus*, which makes the process of exploring and sharing 3D trajectories as easy as browsing a website. We provide a python script that reads trajectory data and enriches them with additional features, such as edge bundling or custom axes and generates an interactive web-based visualisation that can be shared offline and online. The goal of *linus* is to facilitate the collaborative discovery of patterns in complex trajectory data.

## Introduction

In biology, we often face large-scale trajectory data from dense spatial pathways, such as the brain connectivity obtained from diffusion MRI imaging (Liu et al., 2020), or tracking data such as cell trajectories or animal trails (Romero-Ferrero et al., 2018). Although this type of data is becoming increasingly prominent in biomedical research (Kwok, 2019; McDole et al., 2018; Wallingford, 2019), exploring, sharing, and communicating patterns in such data are often cumbersome tasks requiring a set of different software that are often complex to install, learn and use. Recently, new tools have become available for efficiently visualising 3D volumetric data (Pietzsch et al., 2015; Royer et al., 2015; Schmid et al., 2019), and some of those allow the user to overlay tracking data to cross-check the quality of the results or to visualise simple predefined features (such as speed or time). However, given the more general-purpose design of such software, these are not ideal solutions to efficiently and collaboratively explore and share the visualisations. An interactive, scriptable, and easily shareable visualisation (Shneiderman 1996) would open up novel ways of communicating and discussing experimental results and findings (Callaway 2016). The analysis of complex and large-scale trajectory data and the creation and testing of hypotheses could then be done collaboratively. Importantly, since such bioinformatics tools would be right at the interface of computational and life sciences, they need to be accessible and usable for scientists with little or no background in programming. Ideally, the data should be bundled with a guided, interactive presentation in one concise visualisation packet that can be passed to a collaborator. To address these challenges, we have developed our visualisation tool *linus*, making it easier to explore 3D trajectory data from any device without a local installation of specialised software. *linus* creates interactive visualisation packets that can be explored in a web browser, while keeping data presentation straightforward and shareable, both offline and online (Fig 1a). We began to develop this tool when we struggled to find adequate software to explore cell trajectories during zebrafish gastrulation from large-scale fluorescence microscopy datasets (Shah et al., 2019). *linus* allowed us now to interactively visualise and analyse the tracks of around 11.000 cells (starting number) as they moved across the zebrafish embryo throughout 16 hrs. More importantly, it enabled us to share and discuss visualisations with collaborators across disciplines.

**Figure 1.**
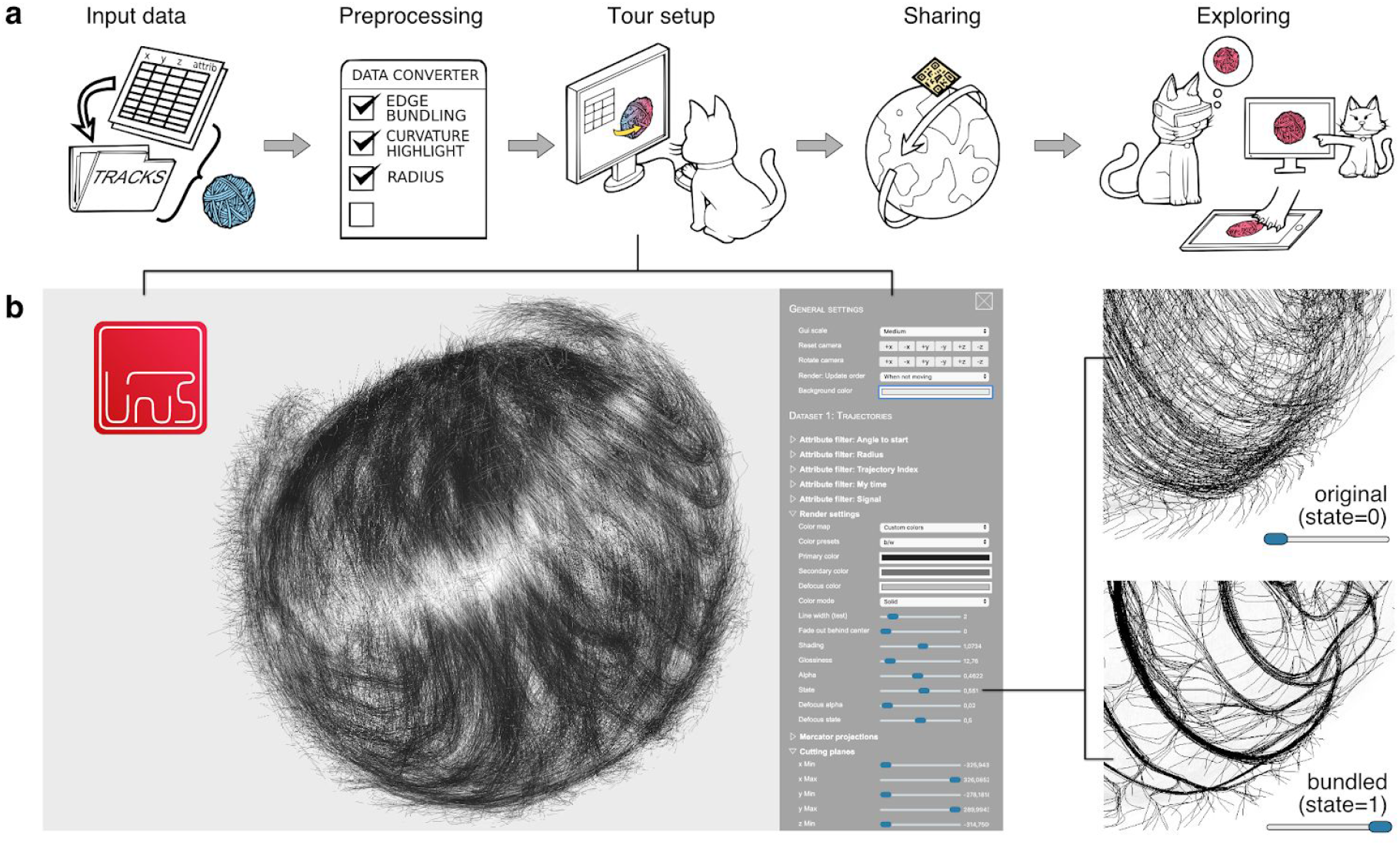
Browser-based exploration and sharing of trajectory visualizations with *linus*. (a) Control workflow of *linus*. Starting with the data, a Python-converter is used to enrich the data with further features (e.g. numeric metrics, an edge-bundled version of the data, visual context) and to prepare the visualisation package. (b) Within minutes, the data can be visualised and explored in the browser, and different aspects of the data can be interactively highlighted (example shows the effect of changing the degree of trajectory bundling).

## Results and Discussion

### linus is a python-based tool that is easy to install and use for scientists at the interface between disciplines

Our overall goal when developing *linus* was to create a versatile and lightweight visualisation tool that runs on a wide range of devices. To this end, we based the visualisation part on web technologies. Specifically, we used TypeScript, which compiles to JavaScript and WebGL. However, a core component of the visualisation process, the data preparation, requires local file access and fast computations, both of which are limited in JavaScript. For that reason, we also created a Python (> v3.0) script that handles the computationally demanding parts of data processing and automatically generates the web-based visualisation packages.

Creating a visualisation package with *linus* is done in a few simple steps (Fig. 1a): The user imports trajectory data from a generic, plain CSV format (see Methods) or from a variety of established trajectory formats such as SVF (McDole et al., 2018), TGMM XML (Amat et al., 2014), or the community standard biotracks (Gonzalez-Beltran et al., 2019), which itself supports import from a wide variety of cell tracking tools such as CellProfiler (McQuin et al., 2018) or TrackMate (Tinevez et al., 2016). During the data conversion, *linus* can enrich the trajectory data with additional attributes or spatial context. For example, we declutter dense trajectories by highlighting the major “highways” through edge-bundling (Fig.1 b). *linus* can automatically add generic attributes that are useful in a range of applications, such as the local angle of the trajectories or a timestamp. The user can simply add custom numerical attributes for specific applications by providing these measurements as extra columns in CSV files (see Methods). The data attributes form the basis for advanced rendering effects. If users want to give a spatial context, *linus* can generate axes automatically, or users can define custom axes.

For more efficient computing, the preprocessing script uses established and optimised packages from python’s rich ecosystem, like NumPy and (Py)OpenCL. In particular, the edge bundling algorithm runs highly parallel on the graphics card and thus, about 10-100 times faster than a CPU-based calculation (with OpenCL-enabled hardware). However, only the creator of a *linus*-based visualisation package needs to run this preprocessor script. The target audience requires only a web browser to view and explore the data. The result of the preprocessing is a ready-to-use visualisation package that can be opened in a web browser on any device with WebGL support. The package is a folder containing HTML, JavaScript, and related files.

### Interactive visualisation with configurable filters allows in-depth data exploration for a variety of applications across sciences

After configuring and creating the visualisation package with the Python toolkit, further adjustments are possible within the web browser. Opening the index.html file starts the visualisation and shows the trajectories with baseline render settings (semi-transparent, single-coloured rendering on a grey background). The browser renders an interactive visualisation of the trajectories and an interface for the user to update and adapt the visualisation to their needs (e.g. colour scales, projections, clipping planes) (Fig. 1b). The user interface itself is adapted to each dataset: The preprocessing script generates a separate property and the corresponding slider (filters and colour mapping) for each given data attribute in the user interface. If more than one state is available for the dataset (e.g. an edge bundled copy of the data, or custom projections), the interface automatically offers the functionality to fade between the states (see Methods).

The user can carve out patterns from the original “hairball” of lines by setting general visualisation parameters like shading and colour maps (Fig. 2a). To focus on particular parts of the dataset, the user filters the data for the various attributes such as specific time intervals or user-specified numerical properties such as marker expression in cell tracking (Fig. 2b). Alternatively, the user can select spatial regions of interest (ROIs) either with cutting planes or with progressively refinable selections (Fig. 2c). The visual attributes can then be separately defined for the selected in-focus areas and the (non-selected) context regions (Fig. 2c) to create a focused visualization. Apart from the purpose of qualitative visualization, the selected trajectories can also be downloaded as CSV files for subsequent quantitative analysis (see Methods).

**Figure 2.**
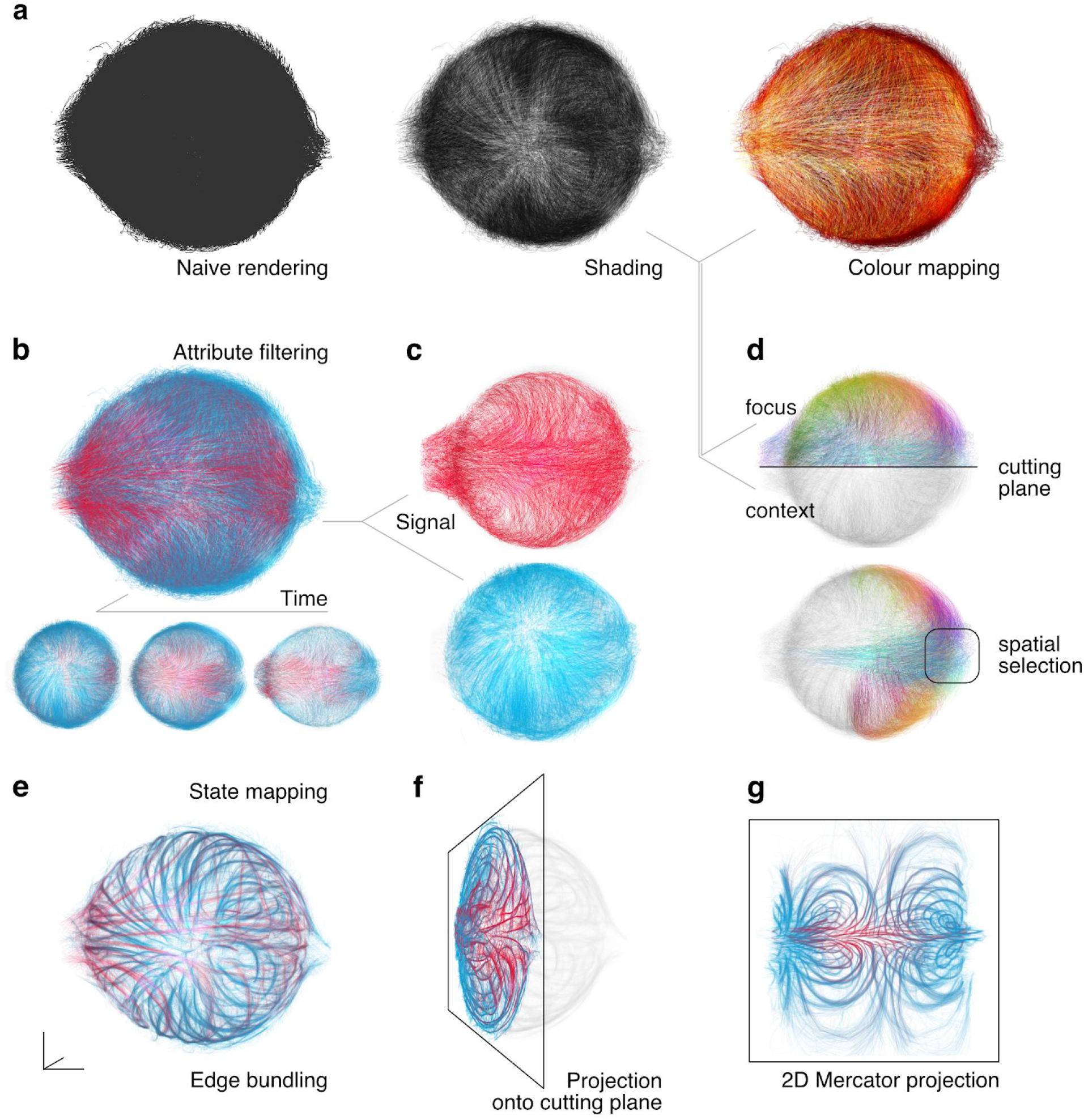
Configurable filters allow deep data exploration. The user can choose from a range of several visualisation methods directly in the browser interface to highlight aspects of interest in the data (zebrafish tracking results from (Shah et al., 2019) as an example). (a) The line data is visualized using a range of options for shading and colour mapping. (b-d) The user can filter parts of the data with respect to specific attributes, such as (b) time intervals or (c) a specific range of signals (marker expression in cells in this case). (d) The user can further create subselections of the tracks in space using cutting planes or refinable spatial selection. The visual attributes can be defined separately for the selected focus region and the non-selected context region. (e-g) The web interface can blend seamlessly between different states of the data. This feature can be used to map between (e) original tracks and their edge-bundled version, to visualize planar projections of the 3D data (f) locally on a definable (oblique) plane or (g) globally using a Mercator projection (with definable parameters).

One important problem with large-scale trajectory data is the sheer density of tracks that often leads to extreme visual clutter. To tackle this problem, one prominent feature of *linus* is the ability to blend between different data transformations seamlessly. We provide two main sorts of transformations out-of-the-box: The user can smoothly transition between original and bundled state to focus on major “highways” (Fig. 2d, Fig. 1b), or between original (3D cartesian) view and different 2D projections (e.g. a Mercator map) to provide a global, less cluttered perspective on the trajectories (Fig. 2e,f). If other, application-specific transformations are needed, such as a spatial transformation or any form of trajectory clustering, the user can provide such an alternative state during preprocessing and then interactively blend between those states.

However, the choice of a web-based visualisation solution brings some drawbacks. The amount of data that can be fluently visualised depends on the underlying hardware (smartphones: >2,000 trajectories, notebooks, and desktop computers: >10,000 trajectories). Another limitation is the reduced feature set which common web browsers offer regarding graphics card access: Compared to the API of OpenGL, the browser-based WebGL API offers fewer shader features. These restrictions lead to some limitations for the rendering process. A drawback of our rendering approach is that it creates artifacts related to the rendering order when we rotate the camera. Thus, we have to order the line fragments *offline* (i.e. not on the graphics card, but in JavaScript), which is a time-consuming process. To maintain high framerates, we only sort line fragments within a second after a user interaction has finished, leading to artifacts during camera motions (see Methods). Furthermore, we cannot provide correct render order when rendering two datasets in the same view, and thus *linus* works best when only rendering one dataset at once.

### Data and visualisations are easily shareable with collaborators via interactive visualisation packets

As a straightforward solution to share the results, the user directly exports the visualisations from the webview as static images and videos (e.g. such as Supplementary Video 1). But sharing the visualisation of the data can go a step beyond image or video data. The user can conveniently record all these visualisation properties directly in the web-interface of *linus* to create information-rich, interactive tours. The user adjusts these tours on a detailed level using a timeline-based editor (Supplementary Fig.3). An icon represents each action that can be moved along the time axis to develop a visual storyline. Smooth transitions and textual markers that can be precisely timed, facilitate understanding and storytelling. To communicate and distribute new findings, these tours can easily be shared online or offline with the community (colleagues, readers of a manuscript, audience of a real or virtual presentation). The tours are copied into the source code of the visualisation package or, if they consist of a limited number of actions (see Methods for details), they can be shared by a dynamically created URL or a QR Code. Fig. 3 shows examples of visualisations that have been created with *linus* ranging from dynamic trajectories in 2D (Fig. 3a) or on surfaces (Fig. 3b) to static (Fig. 3c) or dynamic 3D (Fig. 3d) tracks across applications from ethology, neuroscience, and developmental biology. An interactive version of each example can be found online by simply scanning the respective QR codes in the figure.

**Figure 3.**
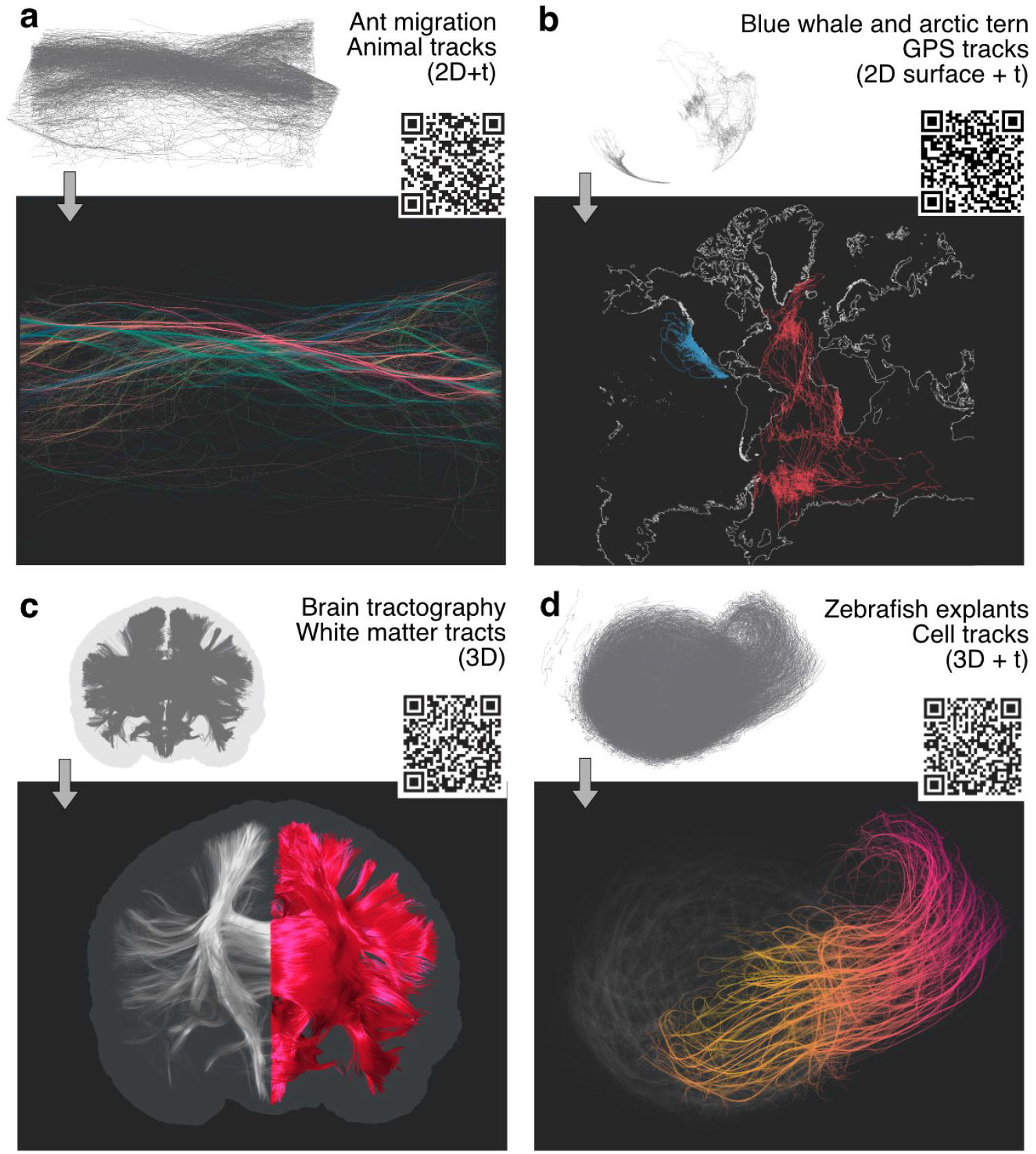
*S*harable interactive visualization packets for a multitude of applications ranging across a variety of sciences. The user can combine the visualization methods, annotations, and camera motion paths in a scheduled tour that can be shared by a custom URL or QR code generated directly in the browser interface. Panels (a)-(d) demonstrate use cases for real-world datasets with different characteristics and dimensionality. (a) Ant trails (2D+t) from (Imirzian et al., 2019). Bundling and colour-coding (spatial orientation by mapping (x,y,z) to (R,G,B) values) indicate the major trails running in opposing directions. (b) GPS Animal tracking data for two species (blue whales (Bailey et al., 2009) - blue and arctic tern (Egevang et al., 2010) - red) shown on a Mercator projection of the earth’s surface. For a better orientation, the outline of the continents is included as axes into the visualization that dynamically adapt to the projections and viewpoint changes (2D surface data + t). (e) Cell movements during the elongation process of zebrafish blastoderm explants (3D+t) (Trivedi et al., 2019). Bundling, colour coding, and spatial selection highlight collective cell movements as the explant starts elongating, focusing on a subpopulation of cells driving this process. Colour code shows time from early (yellow) to late (red) for selected tracks. (f) Brain tractography data showing major white matter connectivity from diffusion MRI (3D). The spatial selection highlights the left hemisphere while anatomical context is provided by the outline of the entire brain (from mesh data) and the defocused tracts of the right hemisphere.

We tested *linus* visualisation packages across various devices and found that performance is the most important aspect of the user experience that varies between different devices. Desktop computers with mid-range graphics cards (e.g. the graphics processors that are built-in with current CPUs) can easily handle more than 10,000 trajectories at smooth framerates. Mid-range smartphones handle the same data with low framerates (ca. 10 fps), which is still usable but does not feel as smooth. For virtual reality applications, we also tested *linus* on the Oculus Go VR goggles. Here, a high frame rate is essential as the user experience would be quite discomforting otherwise and we recommend reducing the number of trajectories further to about 1,000 in this use-case. Due to the differences in performance and user experience, we recommend creating dedicated visualisation packages (or tours) for the intended type of output device.

In the future, we would like to support further advanced preprocessing options such as trajectory clustering, more generic transforms or feature extraction. We also would like to extend the visualization part of *linus*, so the user can interactively annotate the data. Here, we envision that the user can easily label subsets of trajectories and then use this information for downstream analysis (such as building a trajectory classifier).

Our experience with *linus* shows that sharing relatively complex data visualisations in this interactive way makes it much more efficient to collaboratively find patterns in data and to create and discuss figures or videos for presentations and manuscripts. More generally, interactive data sharing is helpful when collaborations, presentations, or teaching occur remotely, as it has been a common situation during the current pandemic. At the same time, during an in-person event such as a talk or poster session at a conference, the target audience can explore the data instantly on their computers, tablets, or smartphones. In any case, touch screens or even virtual reality goggles increase the immersion with more natural controls and true 3D-rendering, helping to grasp the trajectories’ spatial relation. With these features, we are convinced that approaches like *linus* will improve considerably how we collectively explore, communicate, and teach the spatio-temporal patterns from information-rich, multi-dimensional, experimental data.

## Methods

Our software consists conceptually of two parts: a Python-based preprocessing and a web-based visualisation tool. We aimed to move all static and computationally expensive adjustments to the preprocessor, whereas dynamic adjustments to tweak the visualisations are all be performed directly in the web browser later. After running the preprocessor, a folder containing HTML, CSS, and JavaScript files is created (called a visualization packet). These files are opened directly or uploaded to a web server.

### Types of input data

We currently support different trajectory file types directly: TGMM (Amat et al., 2014), biotracks (Gonzalez-Beltran et al., 2019), SVF (McDole et al., 2018), and custom CSV. Most formats are designed to store 3D coordinates plus a timestamp primarily, but no other custom data. However, *linus* supports additional numerical attributes that can then be used to filter or colour the trajectories accordingly. We, therefore, offer a generic CSV format which can be supplemented with custom numerical data: Each CSV file contains the data for a single trajectory, the first three columns represent the coordinates (x, y, z) and any further column is interpreted as another attribute. The columns are delimited by semicolons, and the number of columns must be identical for all CSV files. *linus* reads the first line of a CSV file by default as the header and uses this information to automatically name the respective properties in the user interface. The data converter script then expects a folder that exclusively contains CSV files as input.

### Implementation of data preprocessing

The trajectory data are then converted to a custom JSON format by our python-based preprocessor. Python has the advantage of being executable on a wide range of operating systems and hardware. The preprocessor is used with a command-line interface or by calling the respective commands directly. The command-line interface is easier to use, and it covers the most common cases (e.g. visualising a dataset with custom attributes, and automatically adding an edge-bundled version). For more complex cases, e.g. visualising two datasets at once, or using multiple custom states of the data (e.g. custom projections), users can write their own Python script. We provide detailed and up-to-date documentation in our repository at https://gitlab.com/imb-dev/linus.

Time-consuming operations are implemented using NumPy, and the most demanding process (edge bundling) is handled by an OpenCL script, which increases calculation speed by 10-100 fold. All trajectories are resampled to equal length during the preprocessing step, enabling us to use NumPy’s fast matrix-based algorithms (we use *n* * *m* -matrices, storing *n* trajectories with *m* points in each trajectory). The resulting JSON file then contains a list of datasets. Each dataset holds a set of trajectories that optionally can be further organised into several states, for example, the original data and a projected version. At this point, all data are organized in the same structure as it is required by WebGL (Supplementary Fig. 1), which allows faster loading of the data in the next step.

### Implementation of the web-based tool

The visualisation part runs in web languages (HTML, JavaScript, CSS, WebGL). The JSON file containing the preprocessed data is directly loaded as an object by JavaScript. This part of the software copies the numeric arrays from the JSON file into WebGL’s data buffers like the position buffer, index buffers, and attribute buffers. If a dataset contains more than one state (e.g. an original state and a projected state), these states are stored in additional attribute buffers. Depending on the provided data, we also adjust the shader source code dynamically. For example, we inject variables and specific statements into the shader source code before it is compiled by WebGL. With the dynamic creation of buffers as well as code statements and variables, we pre-build a shader program that is directly tailored to the properties of the respective data. As a result, rendering the data allows quick changes of the visualisation (e.g. color mapping or projections) without the need for updating the datasets on the graphics card, which results in higher frame rates and smooth transitions compared to approaches where data is transformed offline.

In principle, *linus* supports an arbitrary number of attributes and states. However, practically this number is limited by the particular device’s abilities (i.e. its graphics card) and WebGL in general. Typically, we have eight attribute arrays on smartphones and sixteen or more on desktop computers. Our software requires four such attribute arrays for internal purposes, plus one more array for each state or attribute. Thus, for a dataset containing original data, bundled data and two custom attributes (that are shared between the states) we would need eight attribute buffers in total, which can still be managed by a smartphone. Visualising adding additional states or attributes requires devices with more capabilities, like a desktop computer.

### The graphical user interface (GUI)

The user interface (see Fig. 1 and Supplementary Fig. 2) consists of a general part that includes options to change the size of the GUI, the background colour, and camera controls. Furthermore, the user can choose how often the render order should be restored (see section “current technical limitations”). Additionally, several data-specific settings are shown, and this section is further divided into:

- *Filters* for each attribute to only show data within a defined range; if window is a positive value, it will be used to automatically display a range [min, min+window] (while max is ignored).
- *Render settings*, including colour mapping, shading, transparency, which can be independently set for selected and unselected trajectories.
- *Mercator projection* plus rotations that are applied to the 3D positions before the 2D transformation, and mapping the “free” z component to attributes for 2D + feature plots (e.g. space-time trajectories).
- *Cutting planes* can be used to generate a generic 2D projection. Here, the projection plane can be defined by selecting a centre point and a normal direction. Everything above the projection plane is then mapped onto the plane.
- The last part of the GUI offers options to export selected trajectories and also shows a list of available tours. This list is used to start or to load a tour into the tour editor.

### Sharing visualisations and tours

As explained above, the user receives a self-contained package. This package can be opened with any web browser that supports WebGL and can be distributed in multiple ways: It can be locally shared (e.g. sent by email or copied using, e.g. a USB stick) or made easily accessible to a broad audience by uploading it to a web server (as done e.g. on our companion website for this manuscript https://imb-dev.gitlab.io/linus-manuscript/).

The method of sharing the actual visualisation package also influences how an interactive tour can be distributed. In order to make a tour reproducible, they are internally represented by a textual list of actions. This script can be copied directly into the source code of the file main.html of the visualisation package. This method works both for server-based and for file-based distribution of the package. If the visualisation package is hosted on a web server, the tours can also be shared simply with a custom URL and QR code that encodes a tour’s actions. However, the length of such tours is restricted: QR codes are limited in the amount of information they can store, and URLs are usually limited as well (but typically this limit can be configured in the web server’s settings). The commands for camera motion and parameter adjustment (e.g. changing the colour) are concise and only require a few bytes of the URL or QR code. In contrast, textual annotations and especially spatial selections require considerably more space. Thus, sharing a tour by QR codes or URLs usually works for tours without selections and without extensive text annotations.

### Specific considerations for virtual reality devices

The virtual reality mode works only when the visualisation package is hosted on a web server. Further, the way of navigation changes slightly because the head position takes over the task of the camera. For convenience, we introduce the possibility to adjust the height of the dataset and to rotate the data horizontally. Inside the VR environment, no GUI is rendered. To allow controlling the GUI, the user can switch between “2D mode” and “VR mode” instantly.

### Export of trajectories

The user can select trajectories and download this selection. The download may take several minutes as the data must internally be converted into CSV format. The result is a zip folder containing one folder for each data set (usually a single folder), each containing a separate folder for each state of the data (e.g. “original” and “bundled”). Each trajectory is saved as a separate CSV file. It should be noted, however, that the user can only download the resampled trajectories and not trajectories in the raw (temporal or spatial) resolution before the data preprocessing.

### Screenshots and videos

At any time, the user can take screenshots and record videos with the respective buttons in the bottom left corner. Video recording requires an up-to-date Chrome-based browser (Chrome Version 52 or later; other browsers might support it as well but only with enabled experimental features). The output format is WebM, which is currently the only file type that can be directly saved from WebGL.

### Additional technical limitations

In order to offer the tool for a broader range of platforms, we decided to utilise WebGL 1.0. This web standard provides the feature set of OpenGL ES 2.0 (https://www.khronos.org/webgl/), which is limited compared to regular OpenGL versions. WebGL 1.0 is implemented by a wide range of browsers, such as Chrome Version 9, Firefox 4.0, Safari 8.0, iOS 8, Chrome mobile 30 (or newer, respectively).

When rendering a scene containing both trajectories and context, our application must render two different types of geometric primitives (lines and triangles) simultaneously. This can only be performed by two consecutive draw calls: the program first renders all triangles, and then we subsequently render the line segments. Since we need to support transparent rendering, we cannot rely on the z-buffer for determining the spatial order of the segments as this works only for non-transparent geometries (The z-buffer usually tells us if a segment should be drawn or not by checking if already another closer segment has been drawn that would cover the new segment). Thus, we use an alternative to the z-buffer: we sort the geometry first and render it starting with the most distant element. Step by step, we draw elements that are closer to the observer over more distant ones ensuring the correct depth ordering of elements. However, we cannot use this idea to compute the overlap between the set of triangles and the set of line segments since they are different types of primitives and as such, require separate draw calls. As WebGL currently does not have a geometry shader, we cannot mix triangles and lines in one draw call. A consequence is that context can only be rendered as a background silhouette.

Our internal resorting procedure can require a noticeable amount of time (e.g. around 0.5 s for 10.000 trajectories). To ensure a fluent user experience, we use an adaptive strategy and only sort the data when the user stops moving the camera. This can lead to some visual artifacts during the rotation of the camera, but after stopping the motion, the correct rendering order is established quickly. For huge amounts of data, or for devices with low CPU performance (the sorting happens on the CPU, not on the GPU), it is also possible to completely disable the sorting. In that case, we shuffle the rendering order, which at least avoids distracting global patterns introduced by these artifacts.

### Data availability

Exemplary visualizations are available by scanning the QR codes in Fig.1 directly or by visiting

https://imb-dev.gitlab.io/linus-manuscript/

### Code availability

The *linus* software including source code and documentation is freely available at our repository at https://gitlab.com/imb-dev/linus.

## Acknowledgments

The authors are grateful to Gopi Shah and Konstantin Thierbach for sharing data and contributing useful feedback. J.W. received funding from the International Max Planck Research School on Neuroscience of Communication: Function, Structure, and Plasticity (Leipzig, Germany; https://imprs-neurocom.mpg.de). K.A. and V.T. acknowledge funding from European Molecular Biology Laboratory (EMBL) Barcelona and Mesoscopic Imaging Facility, EMBL Barcelona for help with imaging.

## Author contributions

N.S., J.H., and I.R. conceived the project. J.W. wrote the software code. M.H. and N.S. supervised the project. N.S. and J.W. wrote the manuscript. K.A. and V.T. generated the dataset on zebrafish blastoderm explants. All authors read, edited, and approved the manuscript.

## Supplementary Figures

**Supplementary Figure 1:**
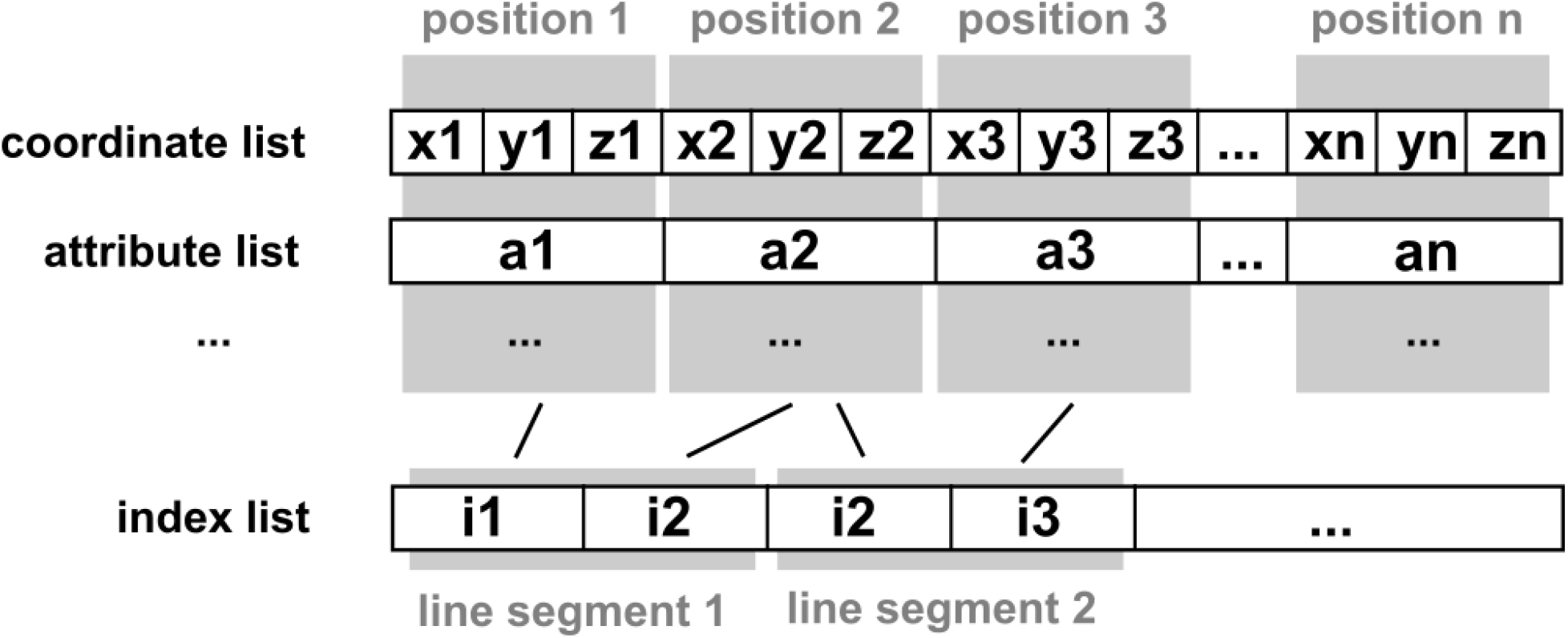
Overview of data structure. The coordinate list holds the x/y/z values for each supporting point of the trajectories. For each such point, an arbitrary number (only limited by the graphics card’s capabilities) of attributes can be stored. The attributes must be provided in the same order as the points. To create trajectories from the point set, an index list is provided as well. Each pair of indices describes one segment of a trajectory. The number of such segments is not restricted, as any point (and its respective attributes) can be used multiple times.

**Supplementary Figure 2:**
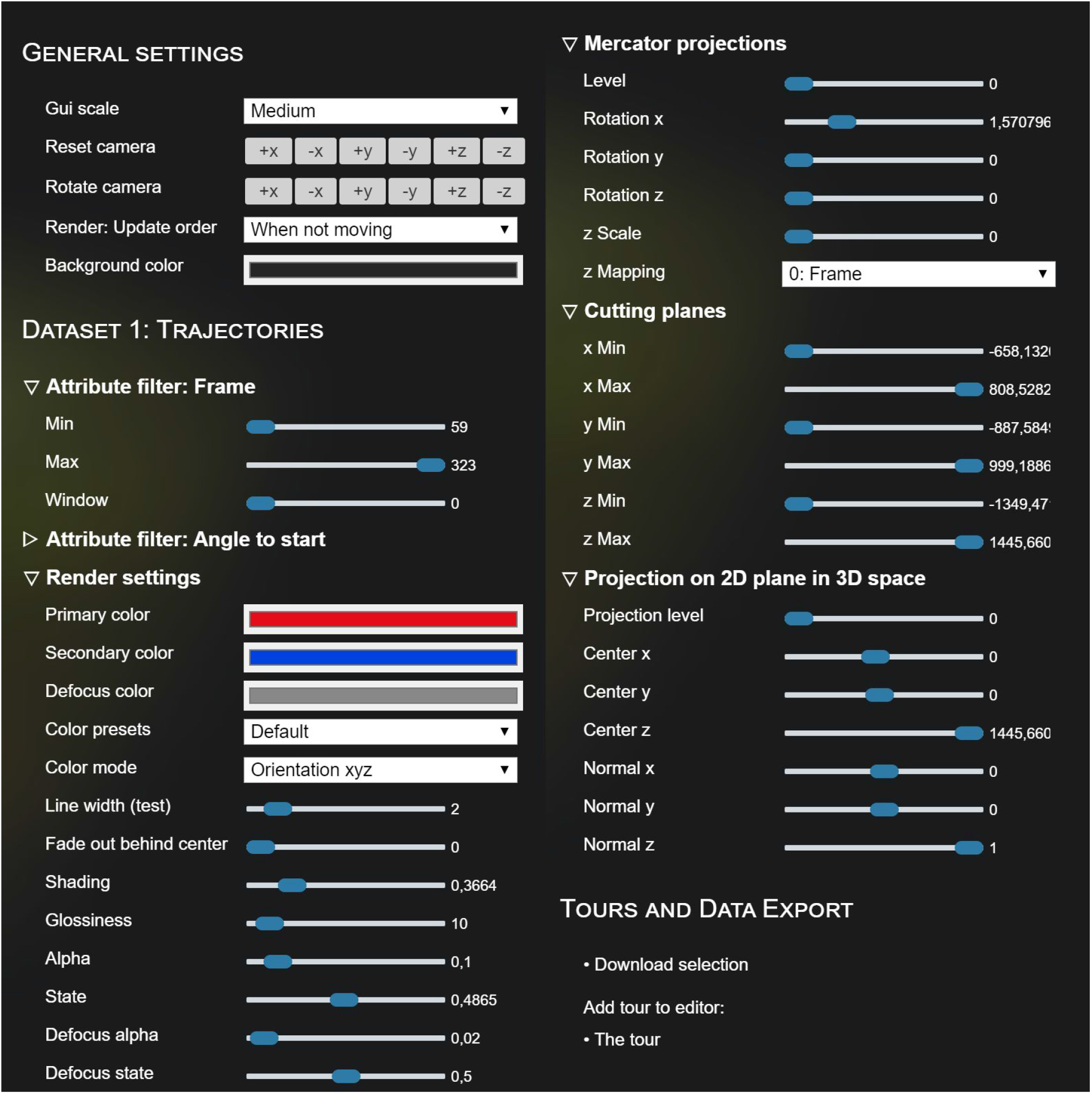
Overview of settings. An overview of the different visualisation settings available to the user from the GUI (two screenshots merged). For explanations regarding different settings, see text or documentation at https://gitlab.com/imb-dev/linus.

**Supplementary Figure 3:**
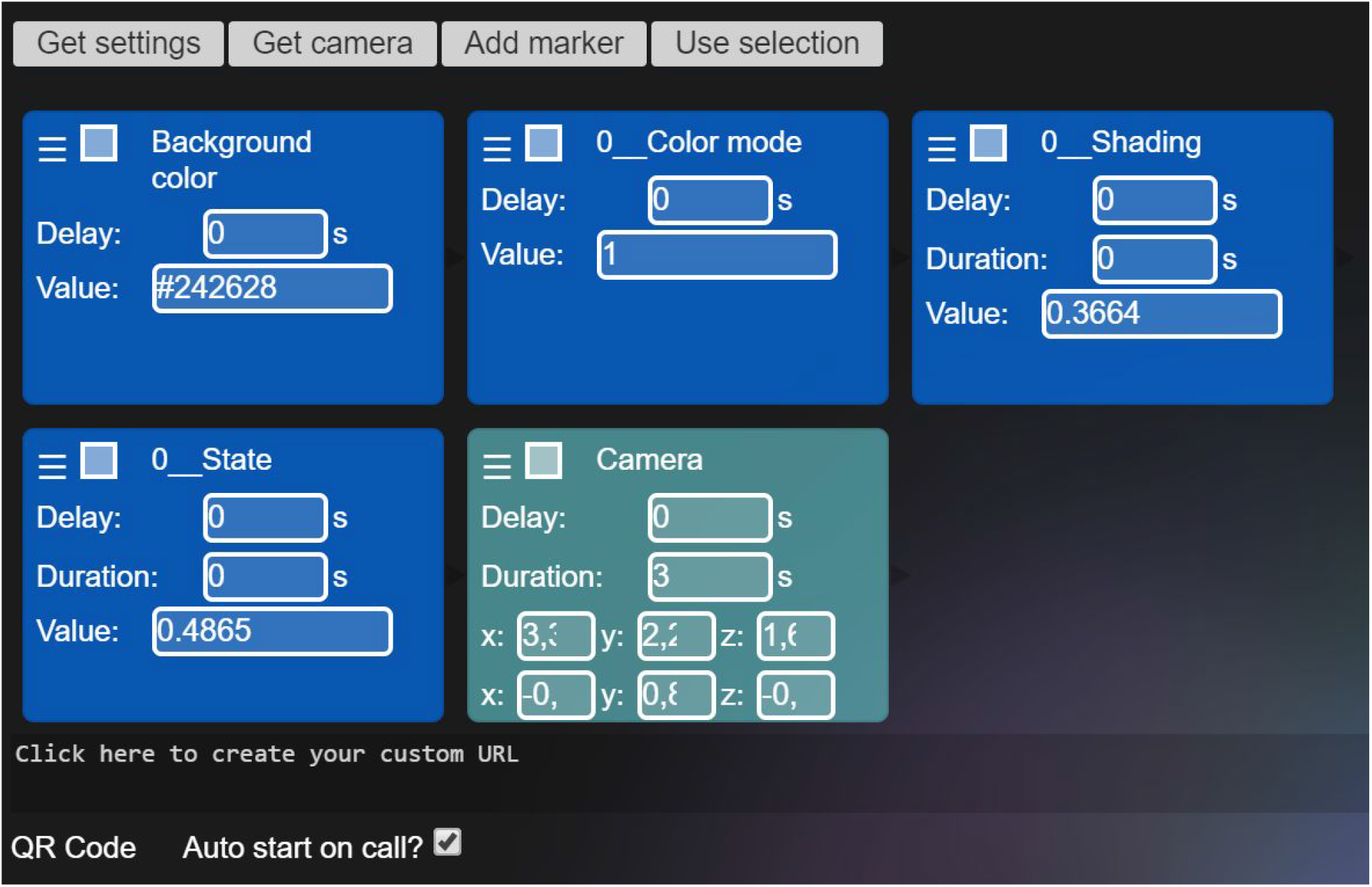
Tour editor. The tour actions can be organised by drag and drop (reading order: from left to right, top to bottom). Every action can be scheduled with a time delay with respect to the end of the previous action. Some actions use transitions (e.g. camera motions or the adjustment of numeric values) whose duration can be configured as well. Eventually, a URL or a QR code can be created.

